# The Promise and Pitfalls of Prophages

**DOI:** 10.1101/2023.04.20.537752

**Authors:** Jody C. McKerral, Bhavya Papudeshi, Laura K. Inglis, Michael J. Roach, Przemyslaw Decewicz, Katelyn McNair, Antoni Luque, Elizabeth A. Dinsdale, Robert A. Edwards

## Abstract

Phages dominate every ecosystem on the planet. While virulent phages sculpt the microbiome by killing their bacterial hosts, temperate phages provide unique growth advantages to their hosts through lysogenic conversion. Many prophages benefit their host, and prophages are responsible for genotypic and phenotypic differences that separate individual microbial strains. However, the microbes also endure a cost to maintain those phages: additional DNA to replicate and proteins to transcribe and translate. We have never quantified those benefits and costs. Here, we analysed over two and a half million prophages from over half a million bacterial genome assemblies. Analysis of the whole dataset and a representative subset of taxonomically diverse bacterial genomes demonstrated that the normalised prophage density was uniform across all bacterial genomes above 2 Mbp. We identified a constant carrying capacity of phage DNA per bacterial DNA. We estimated that each prophage provides cellular services equivalent to approximately 2.4 % of the cell’s energy or 0.9 ATP per bp per hour. We demonstrate analytical, taxonomic, geographic, and temporal disparities in identifying prophages in bacterial genomes that provide novel targets for identifying new phages. We anticipate that the benefits bacteria accrue from the presence of prophages balance the energetics involved in supporting prophages. Furthermore, our data will provide a new framework for identifying phages in environmental datasets, diverse bacterial phyla, and from different locations.

## Introduction

Phages play a central role in microbial dynamics and evolution and have been the source of many of the fundamental discoveries that drove the rise of molecular biology over the last 50 years. Although we comprehensively understand a few phages, most viruses are still uncharacterised (Howard-Varona et al. 2017). The interplay between virulent and temperate lifestyles complicates phage-host relationships. Viruses face the dilemma of choosing one of two disparate life cycles. They can enter a lytic cycle where they infect a host, replicate themselves, and kill their host to release their progeny. Alternatively, they can reproduce as a lysogen integrated into the host’s genome, monitoring cellular health and reverting to lytic growth to exit the host (Lwoff 1953). Temperate lifestyles dominate across many environments where microbes are either highly abundant with high growth rates or in oligotrophic conditions where they are sparse and replicate slowly (Knowles et al. 2016). Once integrated into the genome, temperate phages are vital drivers of bacterial evolution. Lysogenic phages provide novel functions to the bacteria which increase fitness (Edwards, Olsen, and Maloy 2002; Pleška et al. 2018), mediate horizontal gene transfer (Touchon, Moura de Sousa, and Rocha 2017; Sheinman et al. 2021), and protect their host from the assaults of competing phages, plasmids, and other mobile elements through superinfection exclusion (Ebel-Tsipis and Botstein 1971). Identifying prophages remains challenging despite their critical importance to microbial survival and evolution. Prophages have characteristics similar to the surrounding bacterial genome (Akhter, Aziz, and Edwards 2012), the accumulation of transposons degrades the prophage signal (Aziz, Breitbart, and Edwards 2010), and prophages are propagated horizontally (Khan and Wahl 2020). With the rise of metagenome-assembled genomes (MAGs), prophages are often left out of the assembly because their genome composition does not match the rest of the backbone (Papudeshi et al. 2017).

After almost 100 years since the discovery of lysogeny, more than 25 years of microbial genome sequencing, and over a million sequenced microbial genomes, the equivalent characterisation of prophages has lagged, especially across diverse host genomes. The need to improve our knowledge of prophage diversity and the role they play in microbial dynamics has been underscored in the literature for two decades (Casjens 2003; Bobay, Touchon, and Rocha 2014; Touchon, Bernheim, and Rocha 2016; Howard-Varona et al. 2017; Luque and Silveira 2020; Dutilh et al. 2014). Numerous software solutions can identify prophage regions in bacterial genomes, but most utilise homology-based methods to find proteins that look like other phages (Roach et al. 2021). We developed PhiSpy, the first comprehensive approach to identify prophage regions from characteristic genomic features in addition to homology to known prophage genes, and have continuously updated and improved it over the last decade (Akhter, Aziz, and Edwards 2012; McNair et al. 2019).

Here, we use PhiSpy to search all genome assemblies in GenBank – over half a million after quality control – and find that 94 % of those genomes are lysogens containing at least one identifiable prophage. Across most lysogens, we observe a uniform prophage genomic density of 2.4 % (24 bp associated with prophages per 1,000 bp of host genome) and found a massive underrepresentation of prophages in many less studied bacterial taxa. The molecular and biochemical secrets these previously undescribed phages use to infect cells, integrate into genomes, sense their environments, and ultimately kill their hosts are entirely unknown. We will discover many new mechanisms phages use to attack each other, like restriction-modification systems, or bacteria use to protect themselves from phage infections, like CRISPR-Cas, from these new phages.

## Results

We used PhiSpy to identify all the prophages in the publicly available GenBank genome assemblies downloaded on 1^st^ June 2022, as shown in Table 1. We found 540,592 genomes (94 %) that contained at least one prophage. Unless otherwise stated, the following analyses report values from lysogenic genomes.

**Table 1.** Bacterial genome assemblies and prophages identified in our work

|  | Number of genome<br>assemblies | Number of high-<br>quality prophages |
| --- | --- | --- |
| Genome assemblies downloaded | 1,197,396 | - |
| Genomes assemblies processed | 947,091 | 5,007,033 |
| Processed assemblies with fewer than 100 contigs | 574,592 | 2,668,851 |
| GTDB dereplicated genome assemblies | 26,200 | 67,267 |
| Genome assemblies with isolation date | 75,378 | 263,344 |
| Genome assemblies with a country of isolation | 76,070 | 274,148 |

### Lysogens have uniform prophage density

There is a broad correlation between the total bacterial genome length (bp; excluding the prophage sections) and the total prophage length (bp; Fig. 1a) across all the genome assemblies analysed. However, GenBank assemblies are biased towards specific bacterial taxa, especially those associated with human pathogens, because the database contains genomes sequenced for microbial surveillance (Fig. 1b). For example, the food-borne pathogens *Salmonella, Campylobacter_D*, and *Escherichia* comprise 44 %, 7 %, and 4 % of the genomes in the assemblies, respectively (Table S1 lists the abundance of each bacterial genus in the assembly file).

**Figure 1:**
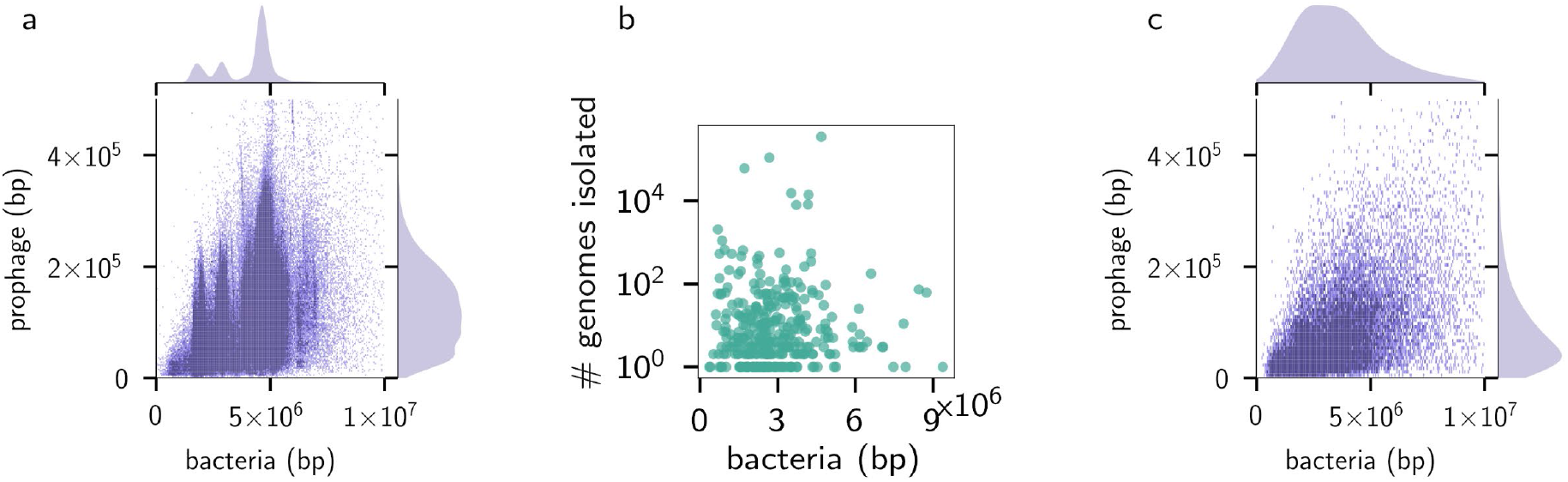
(a) Joint distribution for the prophage and bacterial genome lengths across all 574,609 complete genomes in GenBank. (b) Imbalances in size and the number of genomes isolated (c) Joint distribution for the prophage and bacterial genome sizes per lysogen across taxonomically balanced genomes.

Therefore, we selected a representative set of 26,200 bacterial genomes using the balanced Genome Taxonomy Database (GTDB; version 207) taxonomy (Parks et al. 2020, 2022), incorporating bacterial species proportional to the estimated diversity that each lineage contributes to the entire phylogenetic tree. Dereplicating the genomes by taking one of each species eliminated the sharp peaks in the kernel density estimates (KDE) caused by uneven sampling and revealed unimodal distributions for both bacterial and prophage genome lengths, with maxima at 20.0 kbp for phage DNA and 2.3 Mbp for bacterial DNA (Fig 1c).

We normalised the total prophage length in each genome by the bacterial genome length (to provide bp of prophages per bp of bacterial DNA). We found a uniform distribution across all bacterial genome sizes except for the smallest genomes (those <2 Mbp; Fig 2a). The relationship we found also holds if the number of prophages is normalised by the genome length since there is a strong correlation (R^2^ = 0.82) between the number of prophages and total prophage length (Fig. S1).

**Figure 2:**
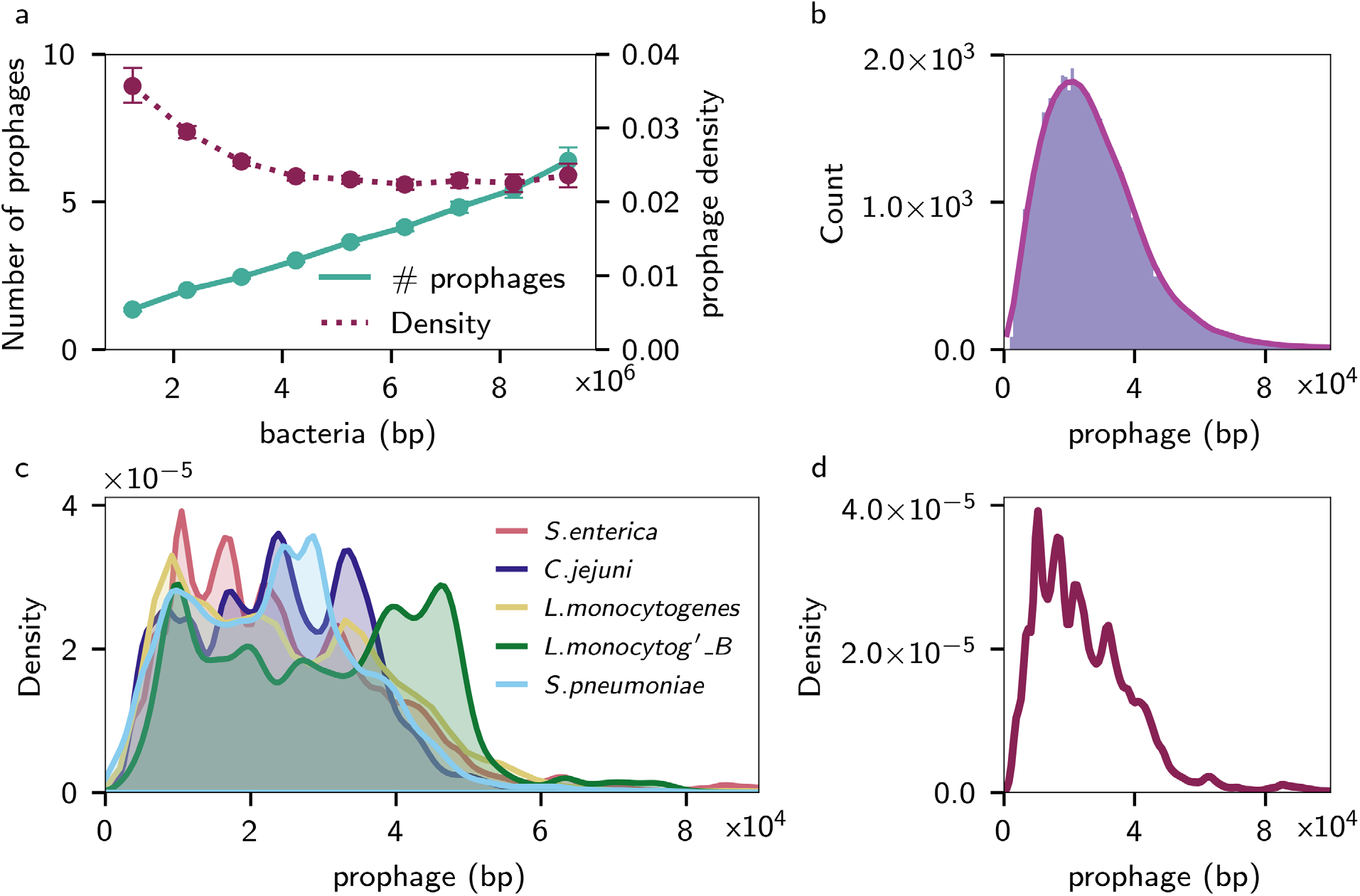
(a) Prophage concentration against base pairs of bacteria, showing the total number of prophages (solid line) or density of base pairs of prophage (dashed) amongst lysogens in a taxonomically dereplicated genome set. (b) The size distribution of individual prophages across the dereplicated GTDB genomes as a histogram (blue bars) and a KDE distribution (red line). (c) KDE distributions for individual prophage sizes across the top 5 most abundant species in the database: from top to bottom in the legend Salmonella Enterica, Campylobacter jejuni, Listeria monocytogenes, Listeria monocytogenes_B, Streptococcus pneumoniae. (d) KDE distributions of individual prophage sizes from 80,000 Salmonella enterica genomes.

Our findings are inconsistent with widely cited analyses that observed increased prophage density with increased bacterial genome size (Touchon, Bernheim, and Rocha 2016). However, we successfully reproduced the observed abundance and density distributions in Touchon *et al*. (2016) by incorporating all the analysed genomes (Fig. S2). An uneven sampling of individual taxa would cause such a discrepancy. Therefore, we explored the distribution of prophages across taxa.

For the prophages predicted from those genomes with fewer than 100 contigs, over half (1,558,270) were longer than 20 kb with a median prophage length of 23 kbp. When only considering the dereplicated GTDB set, the median size was 25 kbp, and the distribution displayed a tailed decay after peaking at 21 kb (Fig 2b).

Prophage sizes across the GTDB set displayed statistically unimodal behaviour (Fig 2b) (Hartigans’ Dip Test, *p*-value 0.99). Our result contrasts the bi- or multimodal prophage genome sizes recently reported in individual or pooled taxonomic groups (Bobay, Touchon, and Rocha 2014; Khan and Wahl 2020). However, those results used limited reference prophages identified from selected bacterial taxa that do not capture the diversity of prophage elements identified in our study.

Nonetheless, within a single species (e.g., *Salmonella enterica*, Fig 2d), the distributions were multimodal; furthermore, multimodal patterns were consistent for many bacteria in the database (Fig 2c, Hartigans’ Dip Test, each *p*-value <1 × 10^−5^). However, there were slightly different locations for the peaks in each taxon (Fig 2c), giving a potential mechanism for the smoothed distribution observed from pooled genomes across diverse and dereplicated taxa

Despite the clear presence of multimodal behaviours within taxa, sampling bias influences the previously observed global prophage distributions, underscoring the methodological challenge of investigating diversity patterns where the data are highly variable and where most genomes represent a small subset of known taxa.

Whilst specific species or genera may have distinct prophage density-genome size distributions amongst taxonomically diverse and dereplicated genomes, the proportion of prophage genetic material in lysogens remains constant at 2.4% for genome sizes above 2 Mbp. Based on that density and the costs of replicating DNA, we estimate that each prophage provides cellular services equivalent to approximately 2.4 % of the cell’s energy or approximately 0.9 ATP per bp per hour.

### Time dependent-trends

We hypothesised there might be temporal patterns in prophage abundances due to the influence of anthropogenic processes such as industrialisation or climate change on the host genomes. We compared the number (Fig 3a) and density (Fig. 3b) of prophages in the genome against the date the genome was isolated as a scatterplot and joint density distribution, respectively. For isolates, we included genomes that explicitly contained isolation date in the metadata, and for metagenome-assembled genomes (MAGs), we assumed the date was the time of sample collection. There were 75,378 genome assemblies with date information and fewer than 100 contigs. Qualitatively consistent results may be found for the entire database, assuming that the sample date is a proxy for the isolation date (Fig. S3). While Fig 3a appears to show increasing prophage abundances over time, linear regression instead indicated a weak trend of fewer prophages more recently (slope -0.02, R^2^=0.007, *p*-value <1 × 10^−121^). However, because we are sequencing more and more genomes over time, and there are more non-lysogens within the joint density distributions (Fig. 3b, hotspot near 2013 and prophage density of zero), we explored the correlation in greater detail.

**Figure 3:**
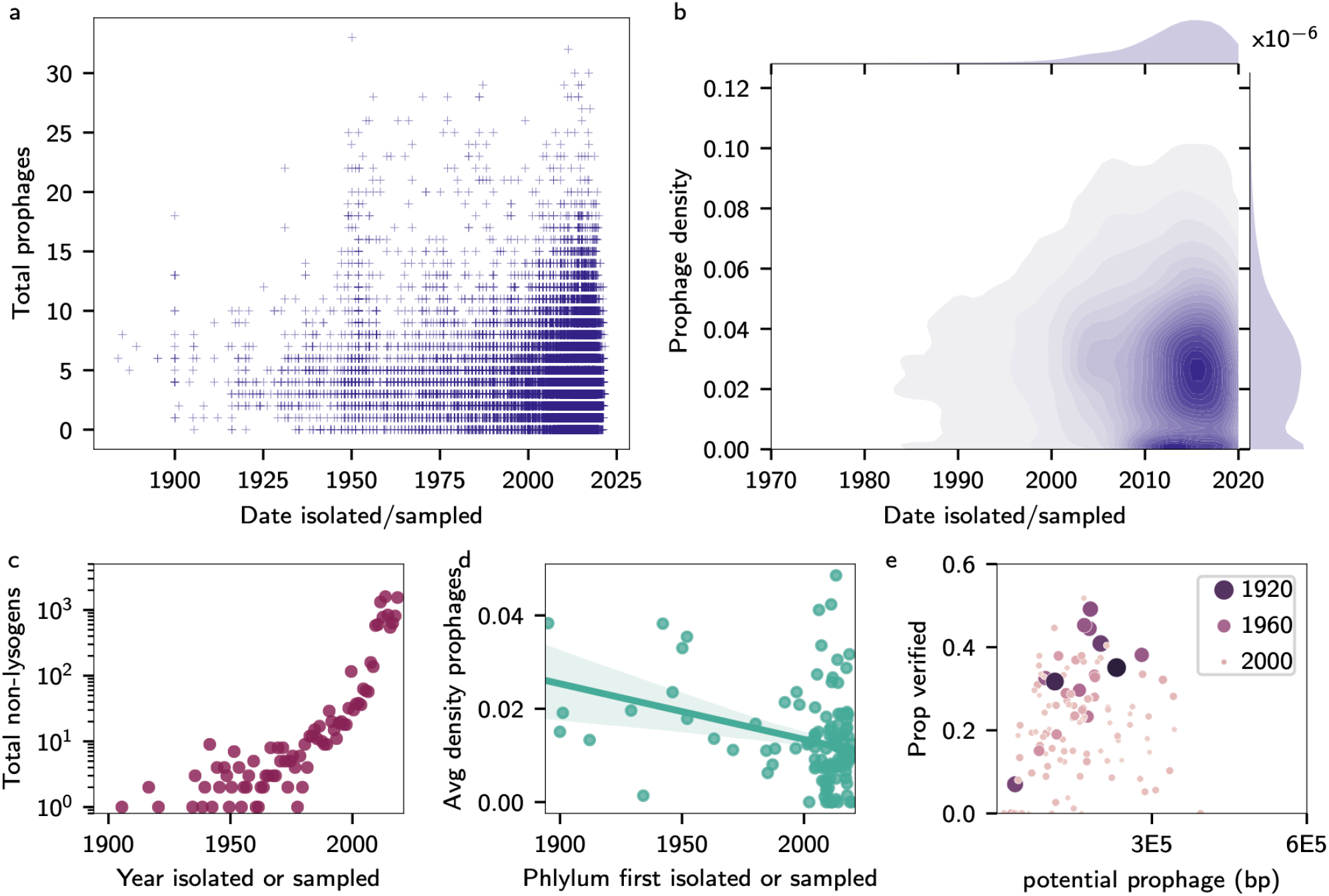
Time trends in prophages. Dates indicate when a genome was isolated or sequenced (for MAGs). (a) Scatterplot of the number of prophages in each genome against the isolation date. (b) Joint KDE for the density of prophages in each genome over time. (c) The total number of genomes that are non-lysogens found over time. (d) A plot of the average density of prophages in a phylum’s genomes compared to the first discovery of an organism from that phylum. (e) Scatterplot of the proportion of DNA with positive hits to the VOG database against the number of base pairs in a genome identified as possible prophage by the machine learning step in PhiSpy. The size and colour of each point correspond to the year the phylum was first isolated, whereby small, light points denote recent events, and large, dark circles denote older events.

Comparing the total number of non-lysogens against the year of isolation shows a sharp jump post-2010 (Fig 3c). The increase in bacteria without prophages was more than the rate of increase seen in lysogens (Fig S4) and corresponds to the advent of low-cost, high-throughput sequencing and the development of metagenome-assembled genome (MAG) pipelines. Most MAG algorithms use a combination of *k*-mer composition and sequencing depth to identify contigs from the same genome. However, prophages do not have the same *k*-mer composition of the bacterial backbone (Akhter, Aziz, and Edwards 2012). In addition, prophages that replicate, even if they do not excise from the genome (Frye et al. 2005), will have altered sequencing depth. MAGs have a 3-fold reduction in mean prophage density within their genomes relative to pure isolates (1.4% and 3.2% respectively, medians 0.73% and 3.0%, Mann-Whitney U test significant at *p*-value <1 × 10^−6^).

Recently discovered taxa are more likely to be described by MAGs, which may imply that less studied bacteria have equivalently less studied phage. To test our hypothesis, we visualised the average prophage density compared to the year each phylum was first described. We found a negative correlation between the time since an organism was first studied and the average density of prophages found in the genome (Fig 3d), a qualitatively consistent relationship at different phylogenetic levels (Fig. S5). We considered whether the negative correlation might result from the under-representation of newly described taxa in the databases or if those genomes may have prophages with qualitatively distinct biological properties instead. PhiSpy identifies prophages through a multi-step algorithm. First, it finds regions that could potentially be prophages because of their unique DNA signatures, including the length of the open reading frames, the number of consecutive open reading frames on the same strand, Shannon score for each open reading frame based on the presence of unique phage *k*-mers, and the deviations of GC and AT skews from the surrounding genome. The software then undertakes a validation step via HMM hits to virus orthologous group databases (VOG). Fig 3e shows the proportion of VOG-verified prophage base pairs against the number of base pairs identified by PhiSpy as potential prophages. The relationship is uniform (linear regression, *p*-value 0.32 and slope <2 × 10^−7^ including several outliers sitting beyond the *x, y* axes limits in 3e), indicating the PhiSpy-proposed viral content of a genome does not correlate to the fraction of hits to viral databases. However, those phyla studied for longer had a higher proportion of verified prophages, whereas recently discovered taxa had more potential prophage sequences discarded, highlighting the presence of substantial database biases across viral and bacterial taxa.

### Geographic and phylogenetic patterns

Significant biases were also evident when examining the taxonomic breadth and depth of sampling across geography. The heatmap in Fig. 4 shows the number of genomes isolated per country (for the 76,070 genomes where country metadata was available). The United States, China, and the United Kingdom had the most sequenced genomes at 25,569, 6,822 and 4,687, respectively.

**Figure 4:**
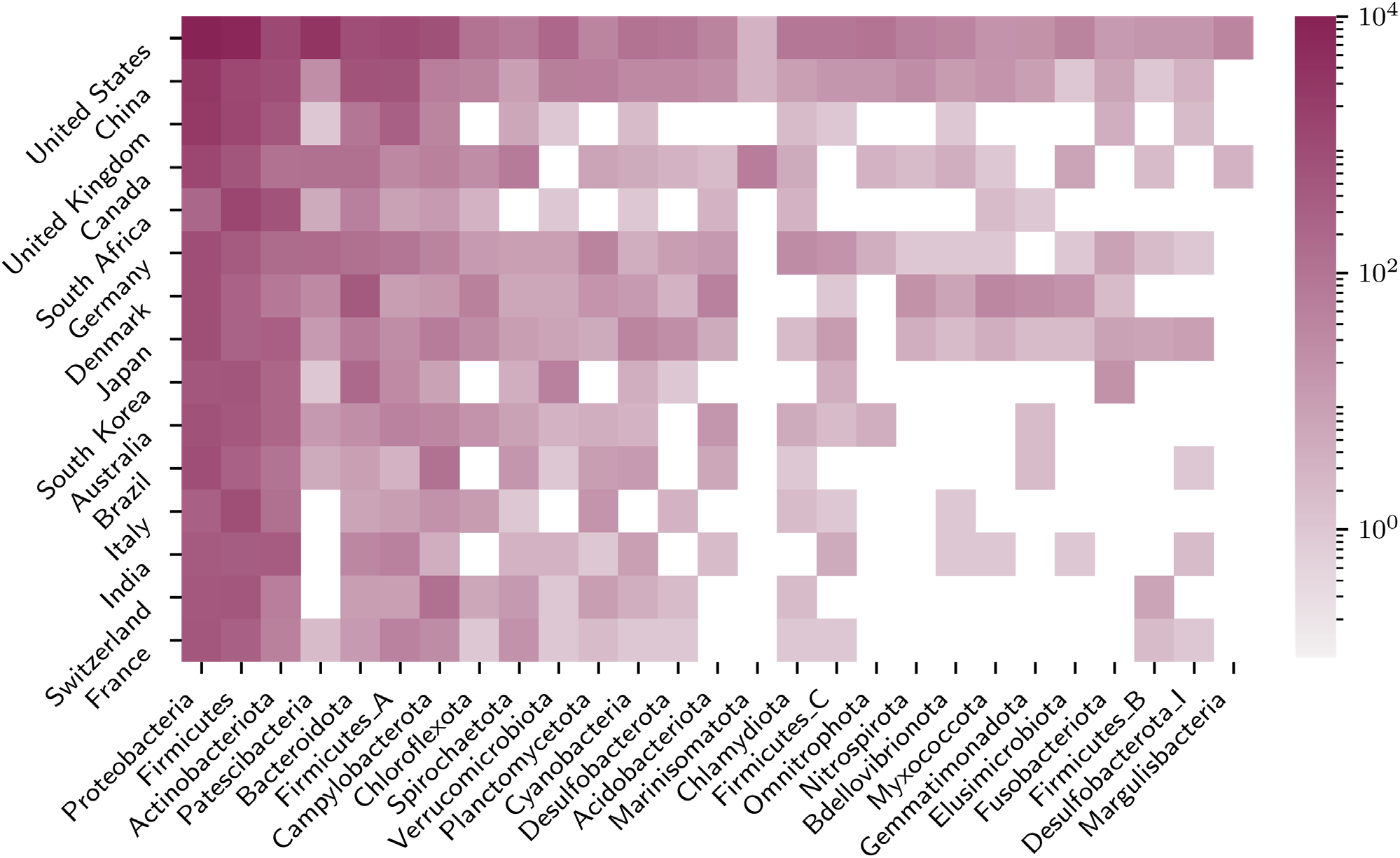
Country and taxonomic signatures for sampling effort. The heatmap shows the top 26 sampled phyla under the GTDB taxonomy and the top 15 contributing countries to Genbank across the 76,070 genomes containing country metadata. The colour scale (log) indicates the number of genome assemblies processed.

There was also taxonomic bias in the isolate genomes: among those genome assemblies with fewer than 100 contigs, the three most sequenced phyla of Pseudomondota (formerly Proteobacteria) (353,392), Bacillota (formerly Firmicutes) (109,590) and Campylobacterota (60,021) contribute 91% of the genomes, primarily due to intensive sequencing of pathogens within those clades. The corresponding proportion of HMM hits of predicted phage base pairs to the VOG database for these phyla were 78%, 55% and 37%, respectively. In contrast, the average per phylum was 30 % across all genome assemblies and 14 % for the five least sequenced phyla with at least five genomes sequenced. Fig. 5 shows the complete taxonomic distribution of prophage densities across the phylogenetic tree. The within-phylum density variance remains consistent at approximately 80% of the mean value.

**Figure 5:**
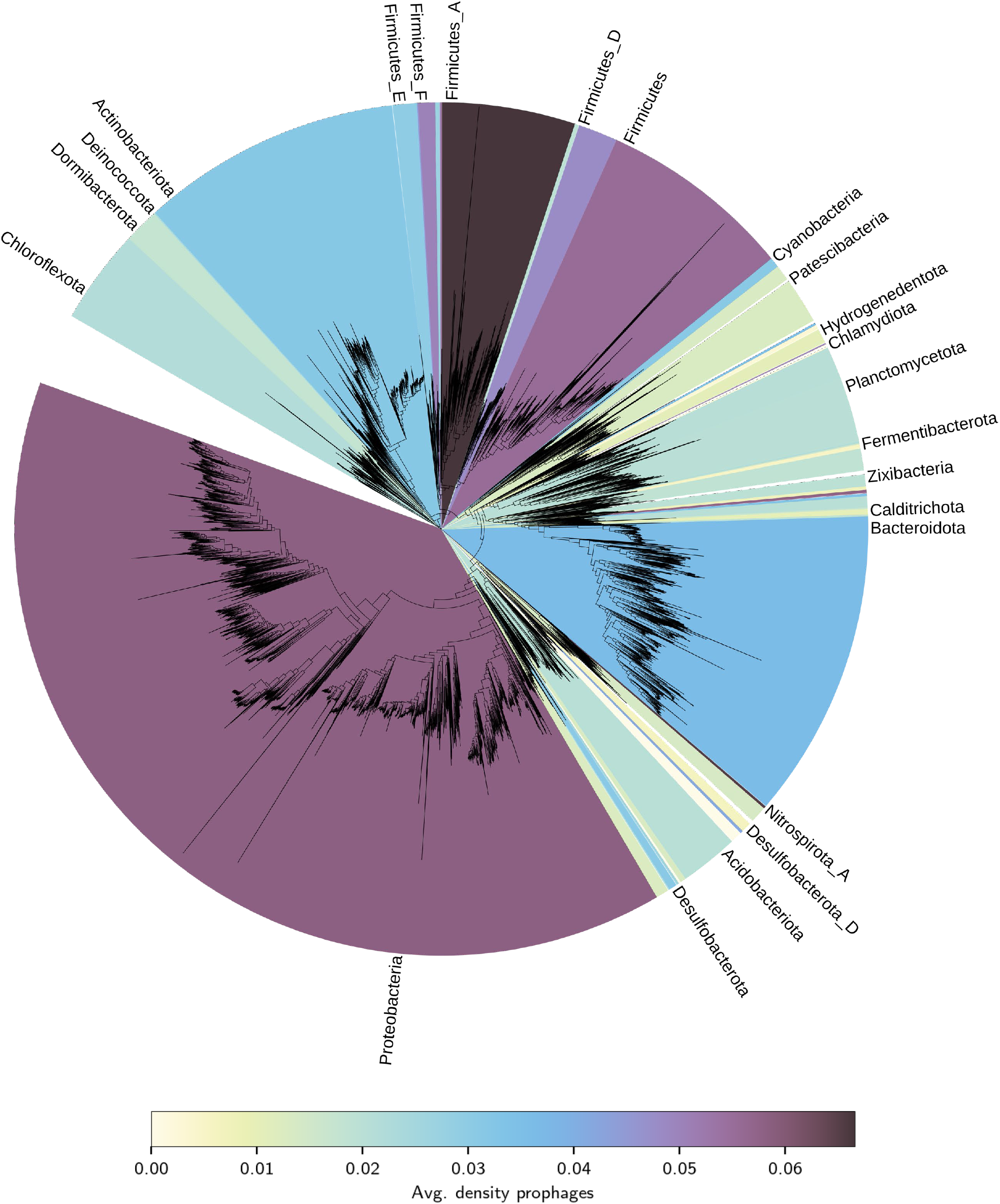
Average prophage density per phylum, visualised across the entire phylogenetic tree from the GTDB taxonomy (Parks et al. 2020, 2022), showing the substantial variation in viral DNA genome content across different phylogenetic groups. The standard deviation of the densities is approximately proportional to the mean density (Fig. S6).

## Discussion

Phages dominate abundance distributions and are critical drivers of bacterial eco-evolutionary processes. The temperate stage of the phage life cycle plays a crucial role in structuring microbial population dynamics through a complex interplay between the life history traits of the virus and its host. Our results provide novel insights into the distribution of prophages across the most diverse set of bacterial genomes to date. We estimate an almost 100% increase in the number of prophages found per genome than previous studies (Touchon, Bernheim, and Rocha 2016) and a higher proportion of lysogens (93%) than previously observed by margins exceeding 10% (Casjens 2003; Touchon, Bernheim, and Rocha 2016; Howard-Varona et al. 2017; Kang et al. 2017; Silveira, Luque, and Rohwer 2021). However, despite the preponderance of lysogens, the density of phage DNA in the genomes was typically lower than the oft-cited upper bounds of 20 % in the literature (Crispim et al. 2018). In contrast to other studies, we found a unimodal distribution of prophage lengths across these genomes, as the central limit theorem predicts, given a large enough stochastic sampling and a finite variance. Our massive sampling showed that the median prophage size was 20 kb.

While our taxon with the most prophage content, *Bartonella tribocorum*, reached an impressive 38.6% prophage density, on average, 3.0 % of the total DNA or 2.4 % for a taxonomically dereplicated genome set was prophage. Touchon *et al*. reported that larger bacterial genomes accumulate more prophages (Touchon, Bernheim, and Rocha 2016). However, that observation was paradoxical because purifying selection for deleting extraneous genetic material is more potent in larger genomes and should select against even short sections of DNA (Bobay and Ochman 2017; Lynch and Marinov 2015). Our findings resolve their conundrum, the number of prophages increases linearly with genome size, but the prophage density across all but the smallest genomes is uniform, indicating a constant carrying capacity of phage DNA per megabase of genomic DNA. There is an optimal balance between the cost of replicating, transcribing, and translating additional prophage genetic material against those prophages’ benefits. Our observation confirms that the benefits prophages confer, especially the protection from other phages through superinfection exclusion, outweigh the evolutionary pressure to reduce genome size (Bobay and Ochman 2017). Across a broad taxonomic range, our best estimate for an evolutionary equilibrium is a prophage density of 2.4 %.

A range of factors affects the prophage density per genome. For example, small genomes, such as those <2 Mb, are typically associated with restricted lifestyles (e.g. obligate intracellular bacteria or obligate endosymbionts) that are often protected from exposure to phages (Casjens 2003; Bobay and Ochman 2017). Across all small genomes (<2 Mb), the average prophage density was 1.6 %, but of those genomes where we could identify at least one prophage, the average prophage density was 2.6%. Although small genomes are more likely to be non-lysogenic, if they carry a prophage, they have a slightly larger average prophage density (2.6%) than larger genomes. We found multiple prophages within taxonomic groups in which lysogeny is extremely rare or (until recently) thought to be non-existent, such as *Chlamidiae* (Touchon, Bernheim, and Rocha 2016; Howard-Varona et al. 2017) or heavily streamlined marine bacteria like *Pelagibacter* sp. (Breitbart et al. 2007; Morris et al. 2020). In the case of *Pelagibacter*, we find multiple examples of prophage elements between 16 and 23 kb in three genomes (*Pelagibacterales* SAG-MED48 (GCA_014653375.1), *Pelagibacter* sp. RS40 (GCA_002101295.1), and *Pelagibacter* sp. RS39 (GCA_002101315.1)).

The selection of organisms for sequencing has introduced significant biases into the datasets, mainly because pathogen source tracking sequences many genomes that end up in the databases. For example, *Salmonella enterica* and *Escherichia coli* had a 3.3 % average prophage density. Pathogens have more prophages because they frequently contribute to disease (López-Leal et al. 2022; Busby, Kristensen, and Koonin 2013). In addition, the PhiSpy algorithm discards many potential phage hits (65.7% across our dereplicated dataset) if they do not meet stringent confidence criteria. These criteria skew validation towards well-studied taxa, especially microbial pathogens. Defective and decaying prophages obfuscate the accurate identification of prophage genomes. However, since our average prophage genome was 20 kb, cryptic prophages did not dominate our data.

Within some phyla, non-pathogenic taxonomic groups may be understudied even if the overall clade appears “well” studied. For example, Proteobacteria and Firmicutes have inflated prophage abundance due to excellent coverage in medically important genera but negligible coverage in other genera. Discovering prophages in environmental samples is likely more challenging than in isolates due to the nature of MAG cross-assembly pipelines, where prophage reads have a lower probability of binning into large contigs (Papudeshi et al. 2017). Searching for these ‘missing’ prophages may be possible by assembling reads that do not otherwise bin or systematically investigating departures between the prophage distributions of environmental and culture-based assemblies within individual strains. Additionally, sampling some environments may induce lysis due to physical impacts such as temperature changes (Tuttle and Buchan 2020), making it difficult to untangle confounding factors that impact our detection of prophages. We provide *all* our predictions, including those we cannot yet validate, so that future genomic analyses can compare them. Our observation of a unimodal prophage size distribution was in stark contrast to almost all prior work (Bobay, Touchon, and Rocha 2014; Crispim et al. 2018; Khan and Wahl 2020). We believe the scale of our analysis, both in the number of prophages—millions of prophages as opposed to several hundred or several thousand— and the diversity of host ranges revealed these differences. We found unique distributions of prophage genome lengths in different taxa, but those distributions combine into a unimodal prophage distribution. We propose a different distribution profile emerges in each environment as the lengths of the prophages from the various taxa in that environment combine.

By examining the geographic metadata, we demonstrated that most samples are medical-based and isolated within particular countries. However, phages from developing countries, un- or under-sampled regions, and biomes will reveal new phage-mediated components to diseases and novel genetic material (Beattie, Lachnit, and Dinsdale 2017; Cazares et al. 2021; Nagel et al. 2016). We propose that global sequencing surveys rectify geographic inequalities (e.g. Edwards et al. 2019) in conjunction with complementary efforts toward sampling to improve coverage of what is unknown rather than known in databases. Indeed, PhiSpy’s unique algorithm provides a potential route to managing one component of the ‘unknown’ sequence problem in phage genomics. Many of the genes identified in the potential (but discarded) prophage sequences by PhiSpy may be genuine, and we encourage the analysis of our data for cross-referencing against metagenome and virome samples. In parallel to the breadth of unexplored viral diversity in metagenomes (Edwards and Rohwer 2005), our results showcase the staggering breadth of prophage diversity across the globe, which we are only beginning to describe. Various approaches across observational and mechanistic experimental studies, statistical analyses, and computational methods are required to unravel the complexity of the lysogenic landscape.

Many questions arise from our data and metadata, which fall outside our study’s scope. For example, the proportion of defective prophages within or across taxonomic groups (Bobay, Touchon, and Rocha 2014; Khan and Wahl 2020), phage host ranges (Rezaei Javan et al. 2019), differences in prophage distributions between laboratory and environmental strains (Pleška et al. 2018), structural organisation of prophages (Brueggemann et al. 2017), or the existence of theoretically proposed but as yet undescribed ‘tiny’ tailed phage (Luque et al. 2020). We have provided all our prophage predictions for future analyses, even those not meeting the stringent criteria used here.

## Methods

### Genomics

The complete genomes were downloaded in GenBank flat-file format (“.gbff”) from NCBIs assembly repository as detailed in the summary file ftp://ftp.ncbi.nlm.nih.gov/genomes/genbank/bacteria/assembly_summary.txt on 1^st^ June 2022 using rsync. PhiSpy (version 4.1.22) was used to identify the prophages in each genome using the parameters described in Table 2. We compared each genome to the VOGDB database (http://vogdb.org/; version 99; downloaded 3^rd^ July 2020) and the PHROG database (version 4; downloaded 1^st^ June 2022). The outputs from PhiSpy were further analysed using Python with the Jupyter Notebooks provided at https://github.com/linsalrob/PhispyAnalysis and https://github.com/jcmckerral/prophage-distributions. The data files are also available in those repositories. All computations were performed on the Flinders University HPC, deepthought (Flinders University 2021).

**Table 2.** PhiSpy Parameters

| Parameter | Value | Meaning |
| --- | --- | --- |
| --number | 5 | To keep a prophage region, it must contain five or more genes |
| --min-contig-len | 5000 | To include a contig in the analysis, it must be longer than 5,000 bp. |
| --phage_genes | 1 | Each region had at least one gene either annotated as a "phage" gene or had a significant hit to the pVOGs database. |
| --extra_dna | 2000 | Look at 2,000 bp on either side of a predicted phage for flanking repeats |
| --metrics |  | The calculation used all ORF metrics (median ORF length, Shannon slope, AT skew, GC skew, and ORFs in the same direction). |
| --min_repeat_len | 10 | Look for flanking repeats 10bp or longer |
| --phmms | VOGs.hmm | Use the pVOGs hidden Markov models |
| --randomforest_trees | 500 | Build 500 random forest trees to classify the data |
| --threads | 2 | Use two threads to analyse the data |
| --training_set |  | Use the generic training set for all genomes |
| --window_size | 30 | Start with a window size of 30 to estimate the phage locations |

The maintenance costs of a cell are estimated to be 0.2 × 10^9^ ATP hr^-1^ for *E. coli* (Lynch and Marinov 2015). Because most of the maintenance cost is replicating DNA, it is similar to other bacterial species. Prophages are ∼2.4% of DNA in cells regardless of the length of the genome, and 0.2 × 10^9^ * 0.024 = 6.6 × 10^6^, meaning that for a 5.1 Mb *E. coli* genome, the ATP cost per base pair per hour is 6.6 × 10^6^ / 5.1 × 10^6^ = 0.94.

### PATRIC metadata

We downloaded the PATRIC (Wattam et al. 2017) metadata from ftp://ftp.patricbrc.org/RELEASE_NOTES/genome_metadata on 1st June 2022. The PhispyAnalysis DateConverter class provided in the GitHub repository https://github.com/linsalrob/PhispyAnalysis, which converts all dates to a unified digital year, cleaned the *isolation date* field. The Jupyter notebooks in that GitHub resource contain the code to correct spelling and typographical errors in the metadata. PATRIC maintains duplicate entries for GenBank assembly accessions if they occur multiple times, and we only retained the first metadata set for each GenBank assembly accession. We merged the metadata with the prophage predictions using the *assembly_accession* field of the PATRIC metadata.

### GTDB taxonomy

We downloaded the GTDB taxonomy summary (version 207) containing taxa metadata for Genbank genomes from https://data.gtdb.ecogenomic.org/releases/release207/207.0/bac120_taxonomy_r207.tsv.gz. We identified missing accessions from the GTDB database. We ran their nucleic acid sequences (*fna files) through GTDB-Tk (version 2.0.0) using the software’s default settings to obtain GTDB taxonomy assignments. We merged the taxonomy metadata with the prophage predictions using the *assembly_accession* field. We randomly sampled all 62,291 unique species from the balanced GTDB taxonomy from all processed genomes containing fewer than 100 contigs. We provide these additional taxonomy assignments in the online material.

### Statistical Analysis

Statistical analysis was performed in Python using the Jupyter notebooks available at https://github.com/linsalrob/PhispyAnalysis and https://github.com/jcmckerral/prophage-distributions. We calculated prophage density (the ratio of total prophage DNA to the bacterial host genome length excluding the prophage region) by dividing the number of base pairs of prophage in the genome by the number of base pairs of host DNA. We also calculated the prophage concentration as the number of prophages per genome. We calculated bins for genome sizes assuming a minimum value of 0, a maximum value of 1.2 × 10^7^, and a step size of 1.4 × 10^6^. Bins for time (years) were annual.

We implemented Hartigans’ dip test (Hartigan and Hartigan 1985) for unimodality using the package ‘diptest’ imported to Python from R (version 0.76-0, https://cran.r-project.org/web/packages/diptest/index.html) with default settings. The species-specific unimodality tests were undertaken on a maximum of 80,000 randomly subsampled genomes for that species, as that was the maximum permitted by the software. We implemented two-sided Linear regression and Mann-Whitney U tests using the scipy package (version 1.6.2).

### Tree visualisation

We downloaded the tree data in Newick format from GTDB (https://data.gtdb.ecogenomic.org/releases/release207/207.0/bac120_r207.tree) and imported it into iTOL (Letunic and Bork 2021). We imported the average prophage density per phylum, and associated colourmap data, as metadata into iTOL. Clades were fully uncollapsed with default settings. iTOL automatically placed phylum labels, and we manually dereplicated them into dominant groups such that text labels did not overlap. The full metadata tables are available at https://github.com/jcmckerral/prophage-distributions.

## Author Contributions

JCM, RAE, BNP, MJR, PD, KMcN, and LKI ran bioinformatics analyses. JCM wrote the paper. AL provided mathematical insights. EAD and RAE conceived the project. JCM, EAD, and RAE wrote the paper with input from all authors.

