## Supplemental Tables and Figures for "The Promise and Pitfalls of Prophages"

#### Supplementary Figures and Tables

*Online Data*

The 20220601 freeze of the data includes PhiSpy run on all genome assemblies available at that time. The data is available on Flinders University's ROADS (FigShare) at https://open.flinders.edu.au/projects/Prophage_predictions/162127. Moreover, each file may also be accessed by its DOI, as shown below.

**Code repositories.**

All the code to repeat this analysis is available in our code repositories at https://github.com/linsalrob/PhispyAnalysis (DOI: 10.5281/zenodo.7749863) and https://github.com/jcmckerral/prophage-distributions.

***Prophage locations* doi: 10.25451/flinders.22299673**

This file is 447M and contains the following information in a tab-separated text file.

| *Column* | *Title* | *Definition* |
|---|---|---|
| **0** | GENOMEID | Genbank genome assembly accession |
| **1** | Contig | Contig ID in the genome assembly |
| **2** | Start | prophage start position |
| **3** | Stop | prophage end position |
| **4** | Length | length of the prophage ((end-start)+1) |
| **5** | #CDS | number of coding sequences in the prophage |
| **6** | Decision | Whether the prophage is predicted or kept. See below. |

The decision field has three options:

i. Too short (<5 genes)

ii. No genes with homology to any known phage gene

iii. Kept means that PhiSpy predicts that it is a prophage region.

***Prophage statistics*** **doi: 10.25451/flinders.22268722**

This file is 18M and contains the following information in a tab-separated text file:

| Column | Title | Definition |
|---|---|---|
| **0** | GENOMEID | Genbank genome assembly accession |
| **1** | Genome Name | Definition of the genome in the GenBank file |
| **2** | Contigs > 5kb | Number of contigs longer than 5 kb (used to predict prophages) |
| **3** | Genome Contigs | Total number of contigs in the genome |
| **4** | Number of Coding Sequences | Total number of coding sequences in the genome |
| **5** | Too short | Number of phage predictions that were too short (less than five genes in the prediction) |
| **6** | Not enough phage hits | Number of phage predictions that did not have a single HMM match to VOGdb version 99 |
| **7** | Kept | Number of high-quality prophage predictions |
| **8** | Note | The outcome of the computation. This column is critical, especially if the sum of prophage predictions is zero |

***Phispy Output*** **doi:** 10.25451/flinders.22317059

This file is 81G and has GenBank format flat files for all the prophage regions we identified. The GenBank files are essentially a slice of all the data from the genome annotation. We added a few PhiSpy-specific outputs, like hits to VOGdb proteins and the E-value of those matches.

Because there is one file for each prophage prediction (5,005,011 files), we separate them into a directory hierarchy and use the NCBI convention for storing files which uses the first three characters and then separates the numbers into directories, so if you have an accession

`GCA_019958295.1`, the GenBank files will be in the subdirectory `phispy/GCA/019/958/295/`

With bash strings, this invocation leads to the directory:

```
BASE=GCA_019958295.1
ls phispy/${BASE:0:3}/${BASE:4:3}/${BASE:7:3}/${BASE:10:3}/
```

***Matches to the PHROGs Database*** **doi:** 10.25451/flinders.22654570

We used MMSEQS2 (version 13.45111) to compare all the protein sequences from prophages to Version 4 of the PHROGs database (https://phrogs.lmge.uca.fr/). The output is tab-separated text with the following columns:

0. Query ID
1. Subject ID
2. Alignment score
3. Sequence identity
4. E value
5. Query Start
6. Query End
7. Query Length
8. Subject Start
9. Subject End
10. Subject Length

***DNA sequences in a fasta file*** **doi:** 10.25451/flinders.22317060

We have extracted all the DNA sequences and placed them in a multiple-sequence fasta file. For each sequence, the sequence identifier is the accession number of the genome (the GENOMEID) followed by the prophage number in that genome. The rest of the header line includes the name of the organism from the GenBank definition line.

***PATRIC Metadata*** **doi: 10.25451/flinders.22299655**

This is a copy of the PATRIC Metadata we used to analyse the bacterial genomes to find prophages. This metadata was downloaded on 1st June 2022. This archive is provided as that is what we used. The latest version can be found at the PATRIC website (http://patricbrc.org/)

***Additional GTDB Taxonomic Classifications*** **doi: 10.25451/flinders.22299658**

We calculated additional GTDB Taxonomies for genomes we analysed for prophages, and that were not yet available in GTDB. These taxonomies were all calculated using r207 of the GTDB and with the gtdb_tk classify workflow (*gtdbtk classify_wf*)

***Genome lengths*** **doi: 10.25451/flinders.22299661**

This 17MB file summarises the bacterial genomes used in the prophage analysis. This file contains the columns GENOMEID, Number of Contigs, Total Length (bp), Shortest Contig (bp), Longest Contig (bp) separated by tabs.

***Assembly summary*** **doi: 10.25451/flinders.22299664**

This 41MB file is an archive of the NCBI Bacterial Genome Assembly Summary file from 1st June 2022 used to predict prophages. The current version of this file is available at ftp://ftp.ncbi.nlm.nih.gov/genomes/genbank/bacteria/assembly_summary.txt, and you can find more details about this file at the NCBI website ftp://ftp.ncbi.nlm.nih.gov/genomes/README_assembly_summary.txt

This version is provided as an archival version reflecting the version we used in our analysis

### Supplementary Tables

Table S1

| Species | Number of genomes | Proportion of database |
|---|---|---|
| Salmonella enterica | 180,389 | 49.2 |
| Campylobacter_D jejuni | 28,638 | 7.81 |
| Listeria monocytogenes | 14,348 | 3.91 |
| Listeria monocytogenes_B | 13,740 | 3.75 |
| Streptococcus pneumoniae | 11,528 | 3.14 |
| Campylobacter_D coli | 9,655 | 2.63 |
| Staphylococcus aureus | 9,275 | 2.53 |
| Escherichia flexneri | 5,587 | 1.52 |
| Klebsiella pneumoniae | 3,950 | 1.08 |
| Escherichia coli | 3,577 | 0.98 |
| Mycobacterium tuberculosis | 2,929 | 0.8 |
| Pseudomonas aeruginosa | 1,985 | 0.54 |
| Streptococcus pyogenes | 1,956 | 0.53 |
| Acinetobacter baumannii | 1,754 | 0.48 |
| Mycobacterium abscessus | 1,497 | 0.41 |
| Clostridioides difficile | 1,395 | 0.38 |
| Escherichia coli_D | 1,261 | 0.34 |
| Enterococcus faecalis | 1,046 | 0.29 |
| Streptococcus agalactiae | 1,006 | 0.27 |
| Salmonella diarizonae | 968 | 0.26 |

### Supplementary Figures

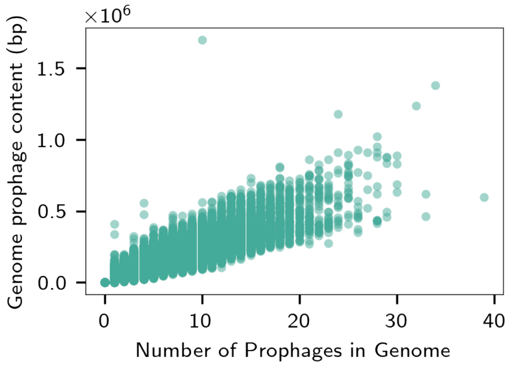


*Figure S1: Correlation between the number of prophages found in a genome and the total number of base pairs of prophage content.*

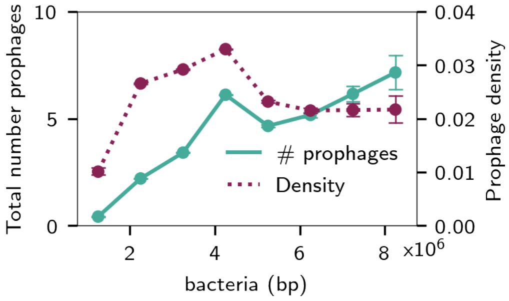


*Figure S2: Prophage total counts and densities across all 574,592 genomes (lysogens and non-lysogens) in the database. This figure reproduces the results of Touchon et al. (2016).*

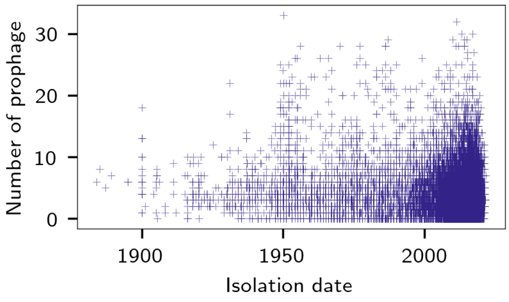


*Figure S3: Prophage counts per genome over time, with the sample date taken as a proxy for the "isolation" date*

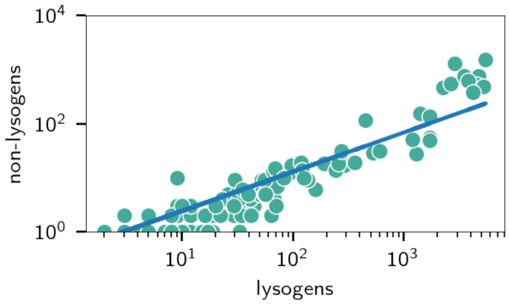

*Figure S4: Each dot represents the yearly number of lysogens and non-lysogens sampled. The total number of genomes (i.e. x coordinate and y coordinate summed together) is a proxy for when the sample was taken (more genomes = more recent). The regression line is fit on data before the structural break (at x ~ 1100) and then projected to the axes limit. There is a proportional increase in the number of non-lysogens once the number of genomes submitted to Genbank in a given year reaches a certain threshold – which corresponds to the advent and rise of MAG submissions to the database.*

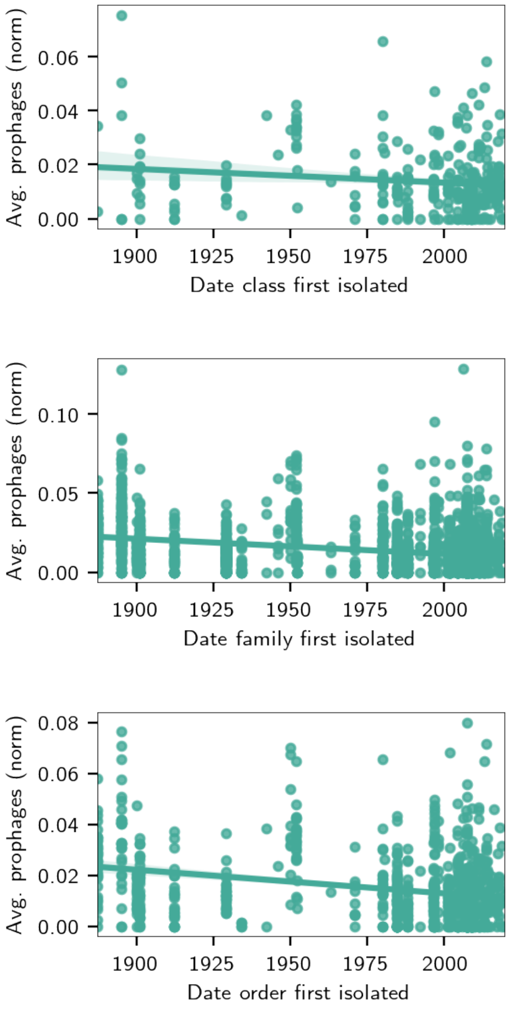


*Figure S5: Average density of prophages in a phylum's genomes, plotted against when an organism from that class (top), order (middle) and genus (bottom) was first sequenced. Taxonomy classifications are from GTDB.*

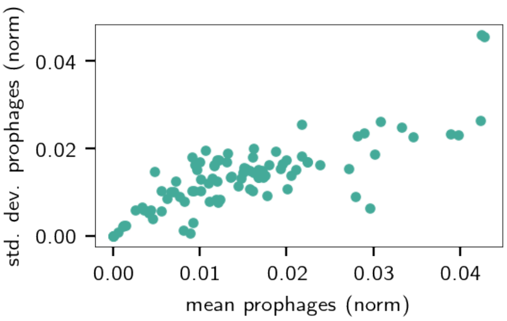


*Figure S6: Standard deviation of mean prophage density (plotted against the prophage density) shows an approximately linear positive correlation between the two.*
